# Proportion and distribution of neurotransmitter-defined cell types in the ventral tegmental area and substantia nigra pars compacta

**DOI:** 10.1101/2024.02.28.582356

**Authors:** William S. Conrad, Lucie Oriol, Grace J. Kollman, Lauren Faget, Thomas S. Hnasko

## Abstract

Most studies on the ventral tegmental area (VTA) and substantia nigra pars compacta (SNc) have focused on dopamine neurons and their role in processes such as motivation, learning, movement, and associated disorders such as addiction and Parkinson’s disease. However there has been increasing attention on other VTA and SNc cell types that release GABA, glutamate, or a combination of neurotransmitters. Yet the relative distributions and proportions of neurotransmitter-defined cell types across VTA and SNc has remained unclear. Here, we used fluorescent in situ hybridization in male and female mice to label VTA and SNc neurons that expressed mRNA encoding the canonical vesicular transporters for dopamine, GABA, or glutamate: vesicular monoamine transporter (VMAT2), vesicular GABA transporter (VGAT), and vesicular glutamate transporter (VGLUT2). Within VTA, we found that no one type was particularly more abundant, instead we observed similar numbers of VMAT2+ (44%), VGAT+ (37%) and VGLUT2+ (41%) neurons. In SNc we found that a slight majority of neurons expressed VMAT2 (54%), fewer were VGAT+ (42%), and VGLUT2+ neurons were least abundant (16%). Moreover, 20% of VTA neurons and 10% of SNc neurons expressed more than one vesicular transporter, including 45% of VGLUT2+ neurons. We also assessed within VTA and SNc subregions and found remarkable heterogeneity in cell-type composition. And by quantifying density across both anterior-posterior and medial-lateral axes we generated heatmaps to visualize the distribution of each cell type. Our data complement recent single-cell RNAseq studies and support a more diverse landscape of neurotransmitter-defined cell types in VTA and SNc than is typically appreciated.

## INTRODUCTION

The ventral tegmental area (VTA) and substantia nigra pars compacta (SNc) are critical structures for processing reward, motivation, learning, and movement. While dopamine (DA) neurons have received the most attention, VTA and SNc neurons that release GABA (γ-aminobutyric acid) or glutamate can mediate related functions and have been implicated in addiction and Parkinson’s disease (Barcomb & Ford, 2023; Buck et al., 2022; Morales & Margolis, 2017). For example, VTA GABA and glutamate neurons contribute to opioid-associated reward behaviors (Corre et al., 2018; Han et al., 2020; McGovern et al., 2023). Also present are neurons that co-express multiple neurotransmitter markers and release multiple neurotransmitters, supported by electrophysiological evidence demonstrating co-release of DA and glutamate, DA and GABA, or GABA and glutamate (Chuhma et al., 2004; Hnasko et al., 2010, 2012; Root et al., 2014; Stuber et al., 2010; Tecuapetla et al., 2010; Tritsch et al., 2012; Yoo et al., 2016).

These neurotransmitter-defined cell types are differentially distributed, some of which are known to be concentrated within subregions defined by cytoarchitecture (Dahlstrom & Fuxe, 1964; Fu et al., 2012). DA neurons tend to be most abundant in lateral substructures of VTA including parabrachial pigmented nucleus (PBP), parainterfascicular nucleus (PIF), and paranigral nucleus (PN). In contrast, VTA GABA neurons are dispersed throughout all subregions, including in posterior regions where the other cell types are less abundant (Ma et al., 2023; Margolis et al., 2012). At the posterior edge of VTA the GABAergic rostromedial tegmental area (RMTg) emerges, however neurons here are genetically distinct from those in VTA and differentially express *Foxp1* and *Pnoc* (Barrot et al., 2012; Jhou, 2005; Perrotti et al., 2005; Smith et al., 2019). Glutamate neurons are most concentrated in medial VTA subregions including interfascicular nucleus (IF) and the rostral and caudal linear nuclei (RLi and CLi) (Kawano et al., 2006; Yamaguchi et al., 2011). While DA neurons are present throughout SNc dorsal, ventral, medial, and lateral subregions (SNcD, SNcV, SNcM, and SNcL), evidence suggests a concentration of glutamate neurons in SNcL (Azcorra et al., 2023; Poulin et al., 2018; Steinkellner et al., 2018).

GABA neurons in SNc and VTA have been known since at least the 1980’s but their distribution remains poorly characterized (Kosaka et al., 1987; Nagai et al., 1983; Oertel & Mugnaini, 1984). Glutamate release from midbrain DA neurons was first demonstrated in primary culture in 1998, and subsequently expression of VGLUT2 was identified in DA and non-DA neurons (Dal Bo et al., 2004; Kawano et al., 2006; Sulzer et al., 1998; Yamaguchi et al., 2007, 2013). However, the literature lacks clear agreement on the relative proportions and distributions of these cell types which have been rarely compared all in the same study (Ma et al., 2023; Nair-Roberts et al., 2008). Here, we used a highly sensitive fluorescent in situ hybridization assay (ISH, RNAscope) to label neurons that express the canonical vesicular transporters for DA (*Slc18a2*, VMAT2), GABA (*Slc32a1*, VGAT), and glutamate (*Slc17a6*, VGLUT2). We selected vesicular transporters as markers because they are required for the vesicular packaging and synaptic release of these neurotransmitters from VTA and SNc neurons. We found that within VTA, DA, GABA, and glutamate neurons were present in similar proportions, though with distinct distribution patterns. In addition, about half of VTA glutamate neurons also expressed VMAT2 or VGAT, in near equal proportion. In SNc, VMAT2^+^ DA neurons represented a bare majority, with GABA neurons also numerous, and a smaller population of glutamate neurons. A subset of SNc DA neurons also expressed either VGLUT2 or VGAT. This quantitative assessment of the relative proportions and distributions of the three major neurotransmitter-defined cell types in VTA and SNc reveals extensive heterogeneity and provides a useful guide for future studies.

## RESULTS

### Triple transporter labeling reveals landscape of neurotransmitter-defined cell types in VTA and SNc

To identify VTA and SNc neurons that release the neurotransmitters DA, GABA, and/or glutamate we used fluorescent ISH (RNAscope) with probes to label mRNA encoding their respective vesicular transporters: VMAT2 (*Slc18a2*), VGAT (*Slc32a1*), and VGLUT2 (*Slc17a6*). The subjects were adult male (n = 3) and female (n = 3) C57Bl/6J mice. For each subject, a selection of 20-micron coronal sections ranging from Bregma -2.5 to -4.3 were labeled by RNAscope plus DAPI nuclear stain and VTA/SNc was imaged by confocal microscopy. Images were assigned Bregma points and brain regions were recognized using anatomical landmarks and by the pattern of vesicular transporter expression. For example, VGLUT2 was prominent in large neurons of the red nucleus (RN) which is devoid of VMAT2 signal; VGAT was abundant in neurons ventrolateral to SNc in the substantia nigra reticulata (SNr) that is devoid of VGLUT2 signal. These together with the pattern of VMAT2 labeling were used to demark VTA and SNc subregions according to Fu & Paxinos (Fu et al., 2012) (**Figure 1, Supplemental Figure 1**). In addition to the signature VMAT2^+^ DA neurons, VGAT^+^ and VGLUT2^+^ neurons were also abundant throughout much of VTA and SNc. While most neurons expressed only one of these vesicular transporters, we also observed neurons expressing all possible combinations of VMAT2, VGAT, and/or VGLUT2 (**Figure 1**, arrows). Importantly, the relative proportion and abundance of these three types varied considerably across VTA and SNc subregions. However, before quantifying their distributions we performed several additional quality control steps.

**Figure 1.**
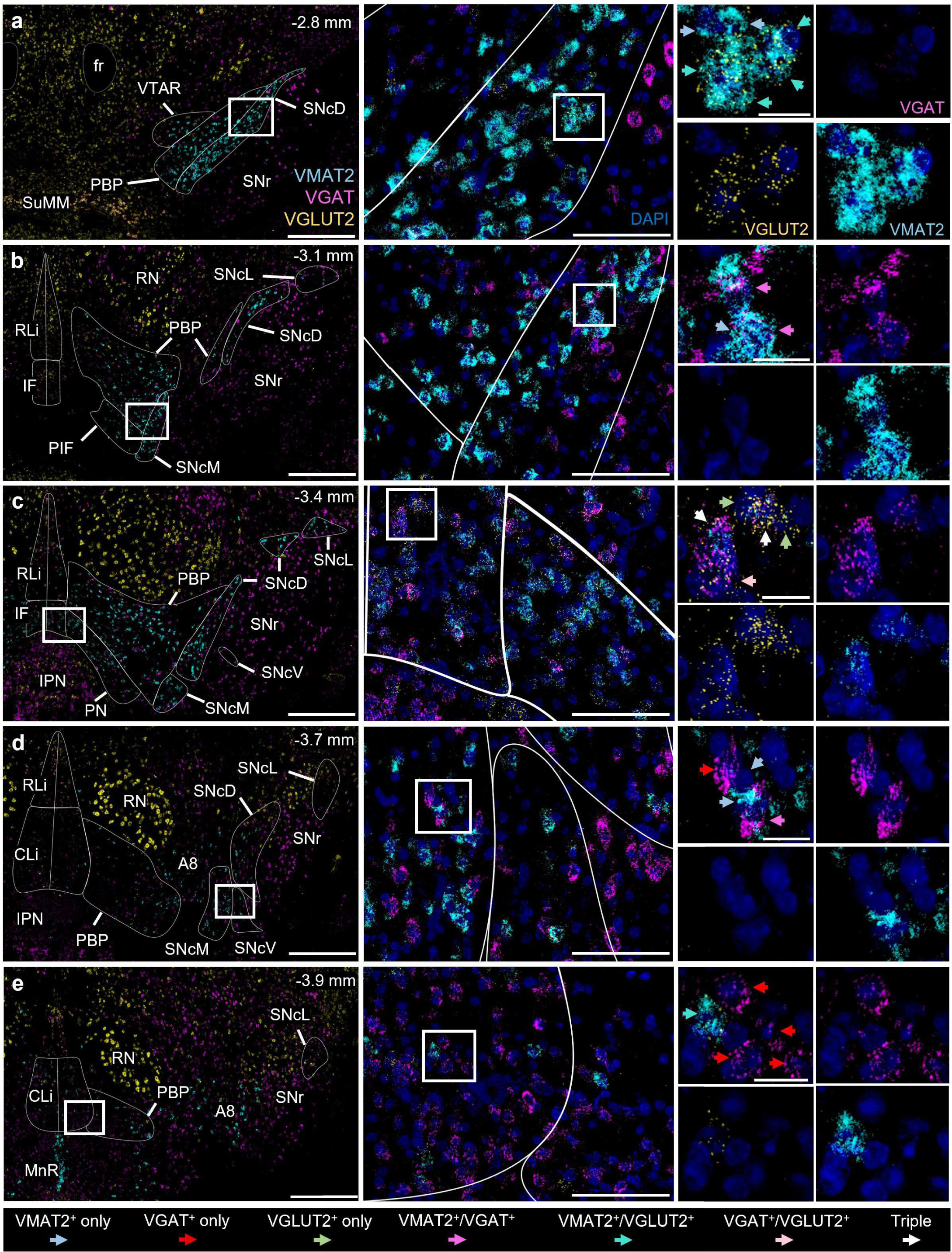
Vesicular transporter mRNA expression in VTA and SNc at various Bregma points. **(A-E)** Example images of coronal sections labeled by RNAscope for VMAT2 (*Slc18a2*, cyan), VGAT (*Slc32a1*, magenta), VGLUT2 (*Slc17a6*, gold), and DAPI nuclear stain (indigo). Left panels show wide-field view with VTA and SNc subregions demarcated, scale 500 µM. Middle panels (scale, 100 µM) represent white box from left panel; right panels (scale, 20 µM) represent white box from middle panel. Right panels show example neurons positive for each of the seven possible combinations of vesicular transporter expression/co-expression, marked by colored arrows. CLi (caudal linear nucleus), fr (fasciculus retroflexus), IF (interfascicular nucleus), IPN (interpeduncular nucleus), MnR (median raphe nucleus), PBP (parabrachial pigmented nucleus), PIF (parainterfascicular nucleus), PN (paranigral nucleus), RLi (rostral linear nucleus), RN (red nucleus), SNcD (SNc dorsal), SNcL (SNc lateral), SNcM (SNc medial), SNcV (SNc ventral), SNr (substantia nigra pars reticulata), SuMM (supramammillary nucleus), VTAR (VTA, rostral).

### Quantification of VMAT2^+^ non-dopamine neurons in caudal-most VTA

Unlike Tyrosine hydroxylase (TH) and other markers commonly used to identify DA neurons, VMAT2 is considered absolutely required for vesicular release of dopamine from midbrain DA neurons (Blakely & Edwards, 2012). However, VMAT2 is expressed not only in DA neurons but also other neurons that package and release monoamines into vesicles. This includes serotonin neurons located in the dorsal raphe (DR), median raphe (MnR), and adjacent structures that overlap with the caudal edge of the midbrain DA cell groups subject to this investigation. Because VMAT2 does not distinguish between these populations of DA and serotonin neurons, we sought to precisely define where VMAT2^+^ serotonin neurons emerge in our samples. To assess the presence of serotonin neurons in caudal VTA subregions, we used RNAscope to label sections (from the same mice as above) for VMAT2 as well as TH and Tryptophan hydroxylase 2 (TPH2), markers of DA and serotonin biosynthesis, respectively.

VMAT2 expression colocalized with TH or TPH2, but never both (**Figure 2A,B**). Sparse TPH2 expression in the caudal VTA began around Bregma -3.6 mm, where they represented 3% of the VMAT2^+^ cells (11/356 cells counted, n = 3 mice) (**Supplemental Table 1**). TPH2^+^/VMAT2^+^ cells were present in increasing frequency in more caudal sections, representing 30% of the VMAT2^+^ cells by Bregma -4.1 mm (8/27 cells counted, n = 2 mice). We estimate that TPH2^+^ neurons, in aggregate, account for approximately 6% of VMAT2^+^ cells in caudal VTA (Bregma - 3.6 to -4.1, 44/702 cells counted, 10 sections, n = 6 mice).

**Figure 2.**
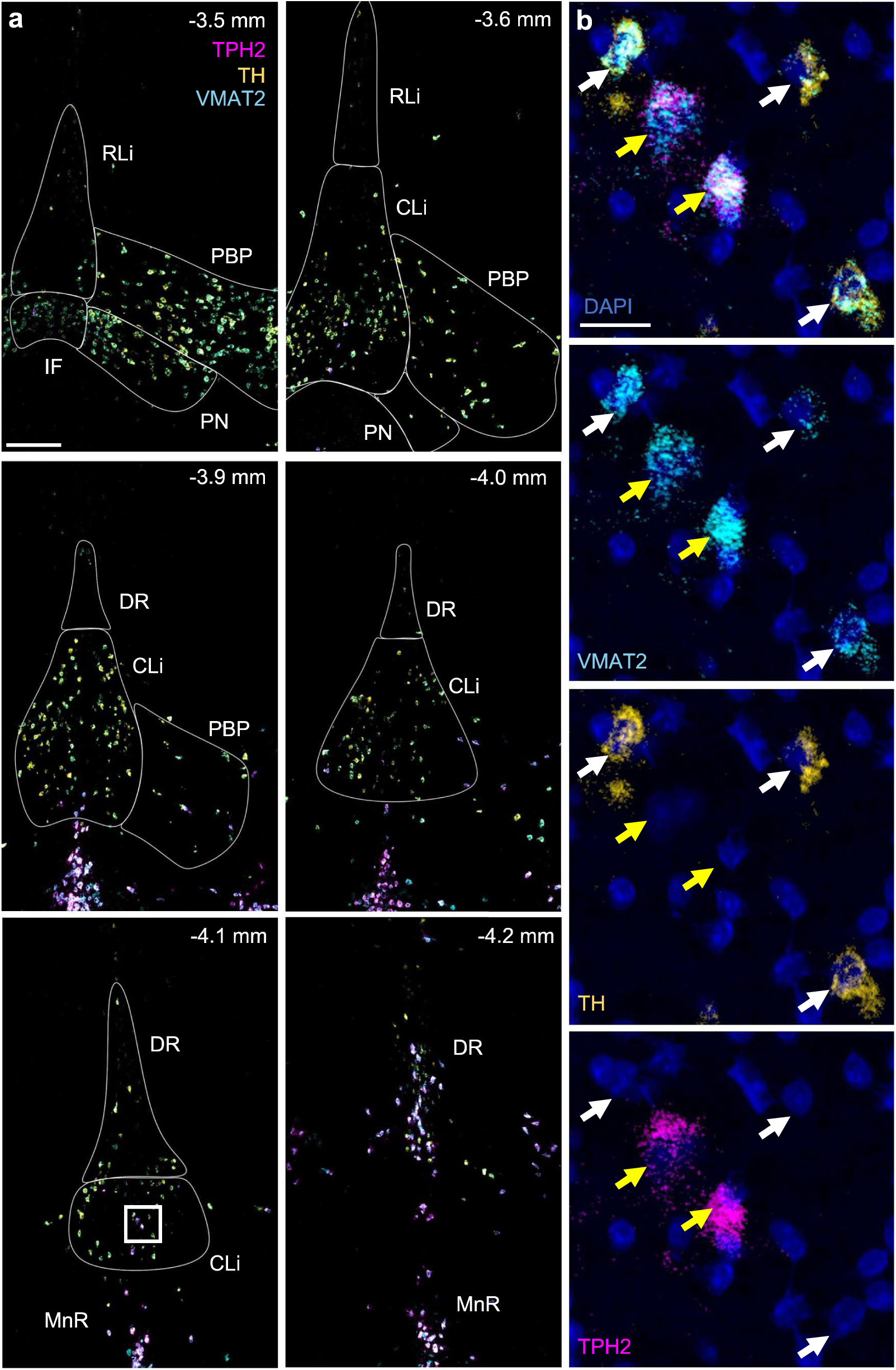
Sparse VMAT2^+^ serotonin neurons in caudal VTA subregions. (**A**): Example images of coronal sections through caudal VTA labeled by RNAscope for VMAT2 (*Slc18a2*, cyan), TH (gold) and TPH2 (magenta). A large majority of VMAT2^+^ neurons express the DA marker TH but not the serotonin marker TPH2. Sparse TPH2 expression in VTA begins around Bregma -3.6 mm and increases caudally. At -4.2 mm Bregma (bottom right panel), CLi becomes indiscernible from DR. Scale, 200 µm. (**B**) Zoom-in of white box from -4.1 mm Bregma shows example neurons dual positive for VMAT2 & TPH2 (yellow arrows), or VMAT2 & TH (white arrows), but no neurons dual positive for TH & TPH2 were detected. Scale, 20 µm; indigo signal shows DAPI nuclear stain. CLi (caudal linear nucleus), DR (dorsal raphe), IF (interfascicular nucleus), MnR (median raphe nucleus), PBP (parabrachial pigmented nucleus), PN (paranigral nucleus), RLi (rostral linear nucleus).

At Bregma -4.2 mm the CLi could no longer be demarcated, in its place remained a population of VMAT2^+^/TH^+^ dopamine neurons localizing to DR. Because DA neurons beyond Bregma -4.1 were heavily intermixed with VMAT2^+^/TPH2^+^ serotonin neurons and localized outside of regions commonly included within the VTA complex, they were excluded from subsequent quantitative analyses.

### Establishing criteria for counting VGLUT2^+^ cells

Cells express variable levels of mRNA for a given transcript, creating challenges when classifying cells as a binary positive or negative. For our study this was a particular challenge for VGLUT2, which tends to be expressed at lower levels within VTA and SNc cells compared to cells in nearby nuclei such as the RN (**Figure 1, 3D**). Preliminary assessments suggested a considerably higher proportion of VGLUT2^+^ neurons within VTA than described in some prior studies. Thus, to first establish levels of background we examined brain regions known to be devoid of VGLUT2^+^ neurons. One such region is the caudate putamen, where we found no discernable VGLUT2 puncta (**Figure 3A-B**). The similar absence of VGLUT2 signal in other regions, such as SNr (**Figure 1**), indicate negligible levels of non-specific background VGLUT2 signal.

**Figure 3.**
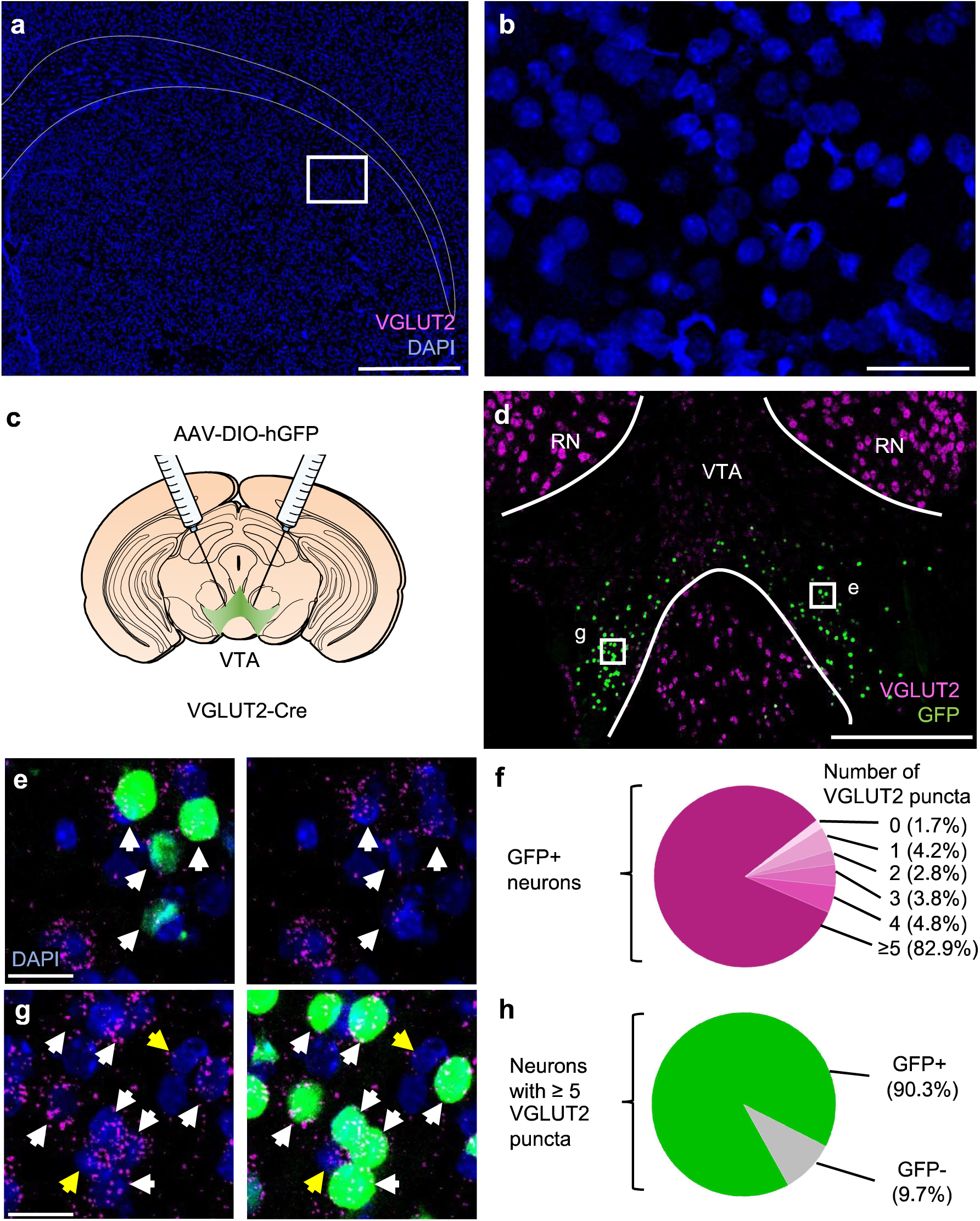
Criteria for classifying neurons as VGLUT2^+^. (**A**) Example images of coronal section through striatum labeled by RNAscope for VGLUT2 (magenta) in caudate putamen. Scale bar 500 µm, indigo shows DAPI nuclear stain. (**B**) Zoom-in of white box from (A) reveals no VGLUT2 signal; scale, 50 µm. (**C**) VGLUT2-Cre mice were injected into VTA with AAV for Cre-dependent expression of histone-tagged GFP. (**D**) Example image of coronal section through VTA immunostained for GFP (green) and labeled by RNAscope for VGLUT2 (magenta); scale, 200 µm. (**E**) Zoom-in of designated box in (D) shows GFP^+^ cells that express ≥5 puncta (white arrows); right image with GFP channel removed. Scale, 20 µm. (**F**) The majority (82.9%) of GFP^+^ neurons contained ≥5 VGLUT2 puncta (n = 1389 cells, n= 3 mice, 10 sections). (**G**) Zoom-in of designated box in (D) containing a high density of GFP^+^ neurons, scale 20 µm. Neurons with ≥5 VGLUT2 puncta were identified as GFP^+^ (white arrows) or GFP^-^ (yellow arrows). (**H**): The majority (90.3%) of identified neurons that had ≥5 puncta also expressed GFP (n = 848 cells, n= 3 mice, 19 sections). RN (red nucleus).

However, when subcellular density of labeling is low it can be difficult to ascribe signal to a specific cell, as is needed for identifying whether more than one mRNA localizes to the same cell. To ensure our threshold for identifying VGLUT2^+^ cells struck a reasonable balance between rates of false positives and false negatives, we cross-validated our approach using VGLUT2-Cre knock-in mice (Vong et al., 2011). AAV for Cre-dependent expression of nuclear-localized histone-tagged green fluorescent protein (GFP) was injected into VTA of male (n = 1) and female (n = 2) VGLUT2-Cre mice (**Figure 3C**), and sections were labeled using a combination of immunofluorescence for GFP and RNAscope for VGLUT2 (**Figure 3D**). To estimate false-negative rate, we counted the number of puncta contained in all GFP^+^ neurons imaged and found 82.9% of these met our criteria of 5 or more puncta (**Figure 3E-F**). This indicates that our criteria excluded up to 17.1% of GFP^+^ neurons that expressed Cre recombinase driven by VGLUT2 regulatory elements. To estimate false-positive rate we first identified GFP-dense areas of VTA, suggestive of high rates of AAV transduction, then identified all VGLUT2^+^ cells using the 5-puncta threshold (n = 848 cells, 3 mice, 19 sections). We found 90.3% were GFP-positive (**Figure 3G-H**), indicating our criteria included up to 9.7% GFP- negative neurons that may not express Cre recombinase driven by VGLUT2 regulatory elements. Because these estimates of false negatives and false positives were similar, we moved forward confident that we were neither grossly under-counting or over-counting VGLUT2^+^ neurons.

### Global proportions of neurotransmitter-defined cell types in VTA and SNc

We next exhaustively counted all cells within one hemisphere of VTA and SNc images that were positive for VMAT2, VGAT, VGLUT2, or a combination of these from each of 48 sections labeled as in **Figure 1** (6 mice, 3F, 3M). For both VTA and SNc, VMAT2^+^ neurons were most prevalent, expressed by 43.5% of 16,789 VTA cells counted, and 53.8% of 8,210 SNc cells counted. Though not powered to detect sex differences, disaggregating these data by sex did not reveal gross differences (**Supplemental Figure 2**).

We were struck by the relative parity with which the three cell types were present in VTA: 43.5% VMAT2, 41.1% VGLUT2, and 37.2% VGAT (**Figure 4B**). These percentages exceed 100% because 20.4% of the labeled VTA cells expressed more than one vesicular transporter. This includes nearly half of all VGLUT2^+^ neurons (45.3%), almost one third of VGAT^+^ neurons (31.4%), and over a quarter of VMAT2^+^ neurons (27.2%) (**Figure 4C**). Of these multi-labeled neurons, VGLUT2^+^/VMAT2^+^ and VGLUT2^+^/VGAT^+^ were most abundant and similar in frequency (**Figure 4A**). For example, 23.2% of VMAT2^+^ neurons expressed VGLUT2 and 24.7% of VGLUT2^+^ neurons expressed VMAT2; while 24.2% of VGLUT2^+^ neurons expressed VGAT and 26.7% of VGAT^+^ neurons expressed VGLUT2 (**Figure 4C**).

**Figure 4:**
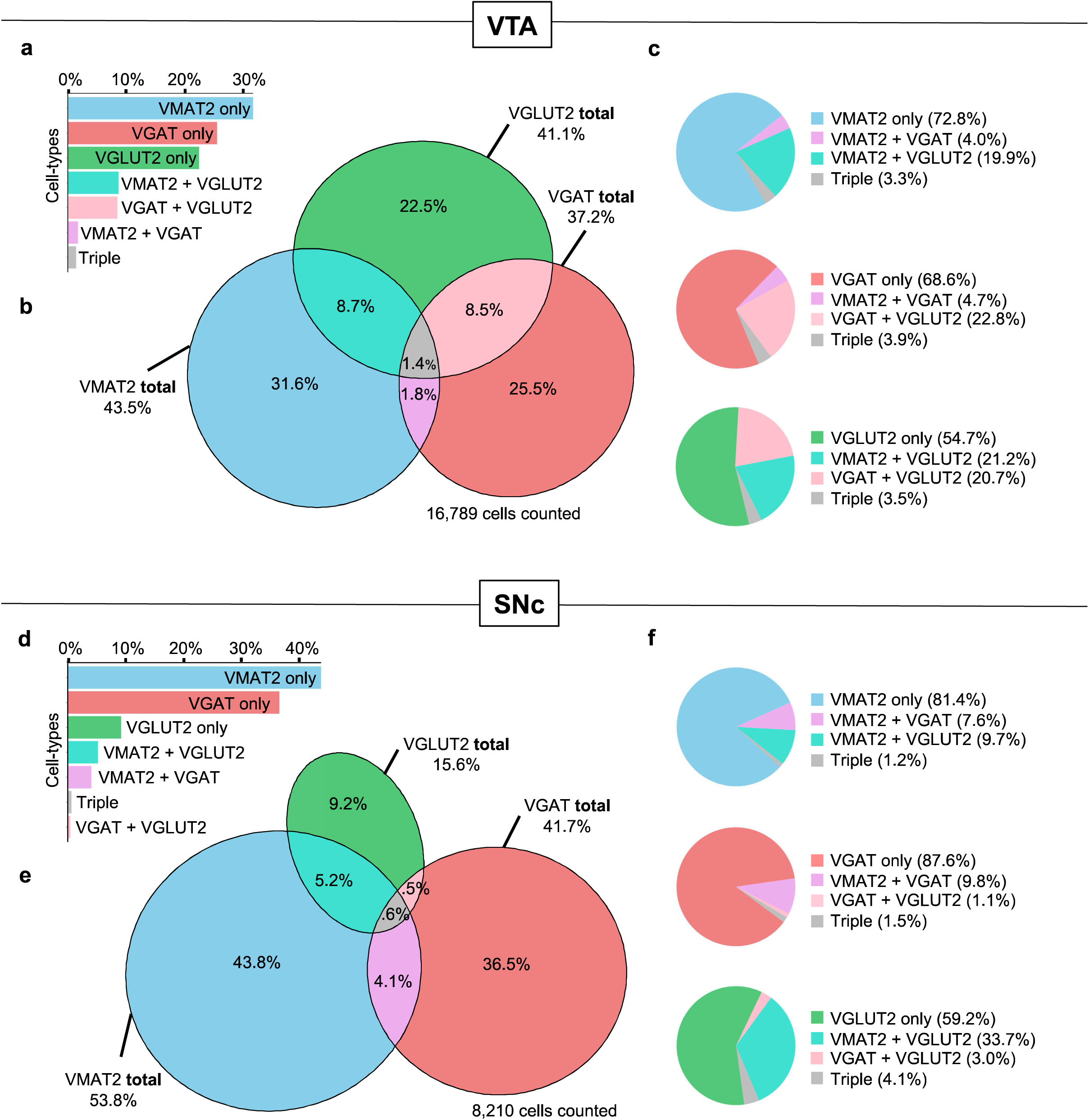
Global proportions of vesicular transporter-defined neurons in VTA and SNc. (**A**) Fraction of labeled VTA neurons that expressed one or more vesicular transporter. (**B**) Venn diagram of vesicular transporter-defined VTA neurons. (**C**) For expression of a given vesicular transporter, fraction of neurons positive for one or more transporters. (**D-F**) Same as in (A-C) but for SNc.

In SNc, 53.8% of neurons expressed VMAT2, 41.7% expressed VGAT, and 15.6% expressed VGLUT2 (**Figure 4E**). Multi-labeled neurons were also present, but about half as frequent as in VTA, representing 10.4% of all labeled SNc neurons. This includes a population of previously identified VMAT2^+^/VGLUT2^+^ neurons, such that 10.9% of VMAT2^+^ neurons expressed VGLUT2 and 37.8% of VGLUT2^+^ neurons expressed VMAT2 (**Figure 4F**).

In addition, we identified a considerable population of VMAT2^+^/VGAT^+^ cells in SNc; 8.8% of VMAT2^+^ neurons expressed VGAT and 11.3% of VGAT neurons expressed VMAT2 (**Figure 4F**). While VMAT2^+^/VGAT^+^ neurons have been observed in single-cell RNAseq and FISH datasets (Phillips et al., 2022; Poulin et al., 2014; Simon et al., 2023), they have received little attention. To cross-validate our RNAscope findings, we crossed VGAT-Cre mice to a ZsGreen (ZsG) reporter line (Madisen et al., 2010; Vong et al., 2011). We immunostained sections from 4 mice (2 F, 2 M) for TH, counted the number of ZsG cells in SNc or VTA from each section, and scored each cell for the presence of TH (**Supplemental Figure 3, Supplemental Table 2**). We found that 14.7% of SNc and 11.7% of VTA ZsG^+^ neurons were co-positive for TH, similar to the ratios identified using RNAscope (**Figure 4**).

### Cell-type composition within VTA and SNc subregion

We next assessed the cell-type composition within each VTA and SNc subregion, guided by boundaries used in Fu and Paxinos (Fu et al., 2012). Within VTA, VMAT2^+^ cells represented the largest share within PIF, PN, and PBP subregions; VGLUT2^+^ neurons the most numerous in RLi, IF, and VTAR; VGAT^+^ neurons the largest share in CLi. (**Figure 5A**). VMAT2^+^/VGLUT2^+^ neurons were most prominent in IF, PN, and CLi; while VGAT^+^/ VGLUT2^+^ neurons were most prominent in VTAR, IF, and RLi.

**Figure 5.**
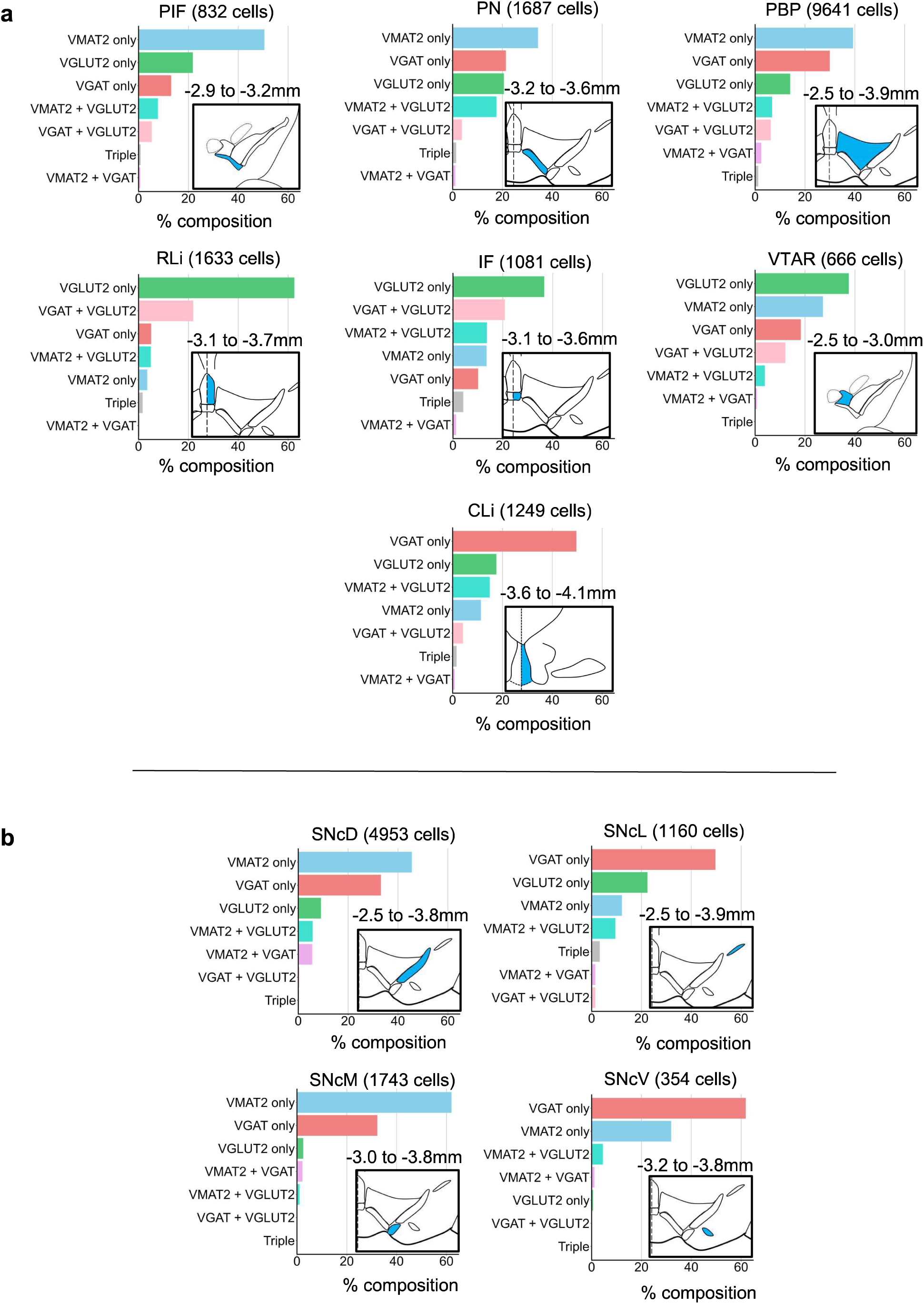
Composition of vesicular transporter-defined neurons across VTA and SNc subregions. (**A**): Fraction of labeled VTA neurons that expressed one or more vesicular transporter by subregion. Inset schematics highlight subregion (blue) with associated bregma coordinates. (**B**) Same as (A) but for SNc subregions. CLi (caudal linear nucleus), IF (interfascicular nucleus), PBP (parabrachial pigmented nucleus), PIF (parainterfascicular nucleus), PN (paranigral nucleus), RLi (rostral nucleus), SNcD (SNc dorsal), SNcL (SNc lateral), SNcM (SNc medial), SNcV (SNc ventral), VTAR (VTA, rostral).

Within SNc (**Figure 5B**), VMAT2^+^ represented the largest share of labeled neurons in SNcD and SNcM subregions. VGAT^+^ were the largest share of labeled neurons in SNcL and SNcV subregions. VGLUT2^+^ neurons, including VGLUT2^+^/VMAT2^+^ neurons, represented a relatively high proportion of cells in SNcL, consistent with prior reports (Poulin et al., 2018; Steinkellner et al., 2018, 2022). VMAT2^+^/VGAT^+^ cells were most prominent in SNcD. Yet, because there were apparent cell-type distributions that transcended these cytoarchitecturally-defined subregions we next analyzed data across anatomical space independent of subregion borders.

### Cell-type distributions across anatomical space

To assess the distribution of VTA and SNc cell types across the anterior-posterior (AP) axis we plotted cell counts by Bregma point. In VTA (**Figure 6A**), VMAT2^+^ neurons were centered within VTA, peaking between Bregma -3.0 to -3.4. VGLUT2^+^ neurons peaked sharply where the midline subregions IF and RLi first appeared at Bregma -3.1, but otherwise show an AP distribution and frequency similar to VMAT2^+^ neurons in this analysis. VGAT^+^ neurons were also widely distributed, but shifted posterior with respect to VMAT2^+^ and VGLUT2^+^ neurons within VTA.

**Figure 6.**
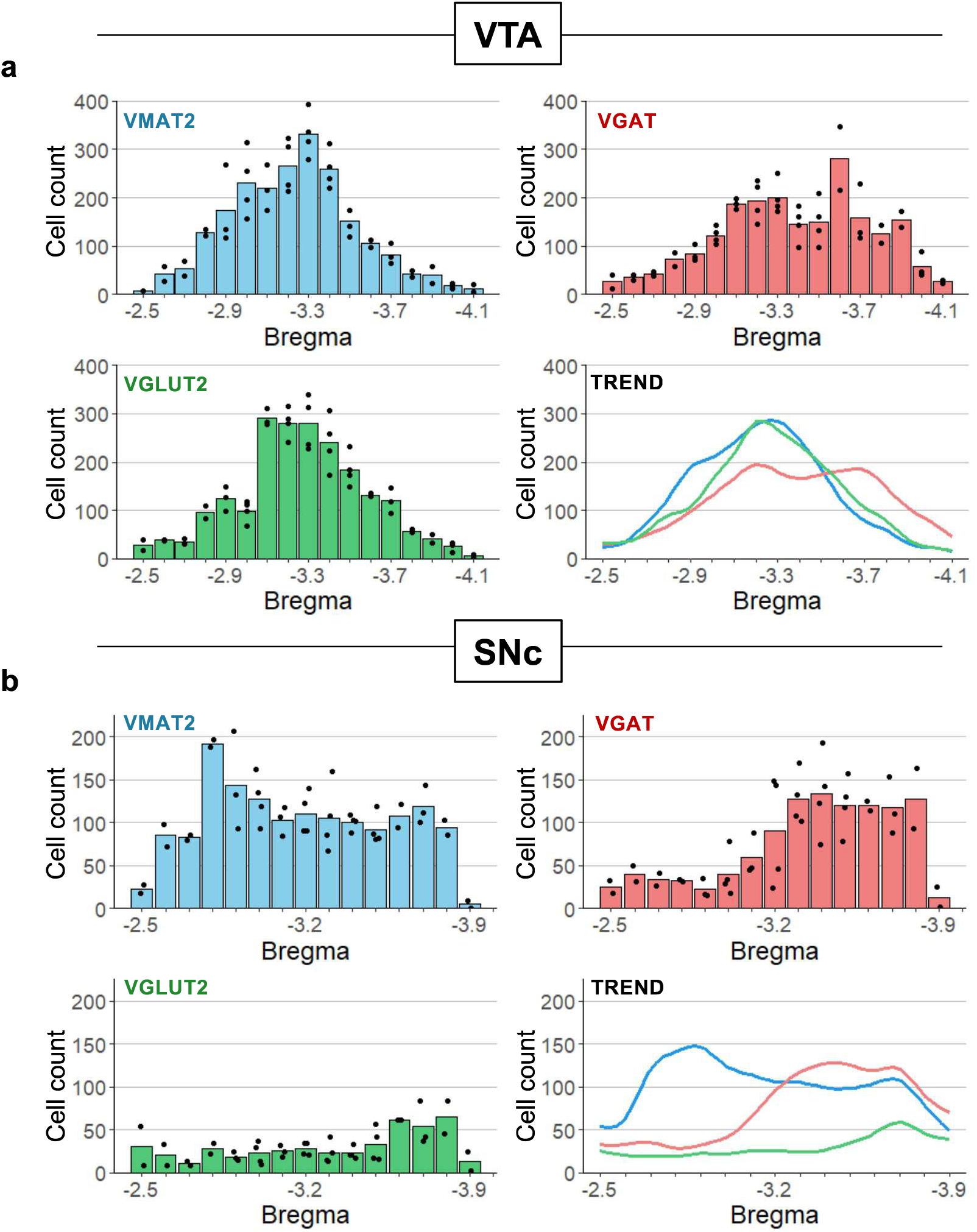
Anterior-posterior distribution of neurons defined by vesicular transporter expression. (**A**) Mean number of neurons expressing VMAT2, VGAT, and VGLUT2 across Bregma point. Trend lines are moving averages smoothed with spline regression. (**B**) Same as (A) but for SNc.

In SNc, VMAT2^+^ neurons peaked at Bregma -2.8 mm and then plateaued (**Figure 6B**). As in VTA, VGAT^+^ neurons were more abundant in posterior sections through SNc. VGLUT2^+^ neurons were present throughout, but at relatively low levels.

The distribution of cell counts presented in **Figure 6** does not account for the changing shape of VTA and SNc across Bregma points (and thus the density of cell types). Nor does it account for the significant heterogeneity across the medial-lateral axis of VTA or SNc. We thus sought to display cell-type density distributions across medial-lateral (ML) and anterior-posterior (AP) space, plotting the equivalent of 100 micron^2^ voxels collapsed across the dorsal-ventral dimension. Some ML-AP coordinates were represented by different numbers of sections, which would bias the distribution plots if total counts were used for each coordinate. To circumvent this, we first calculated the average number of positive cells for each coordinate. We then summed the averages across ML-AP space and plotted these averaged values as a percentage of the summed averages (**Figure 7**). By collapsing the data within each axis and normalizing the result, we also visualized density solely along either ML or AP axis.

**Figure 7.**
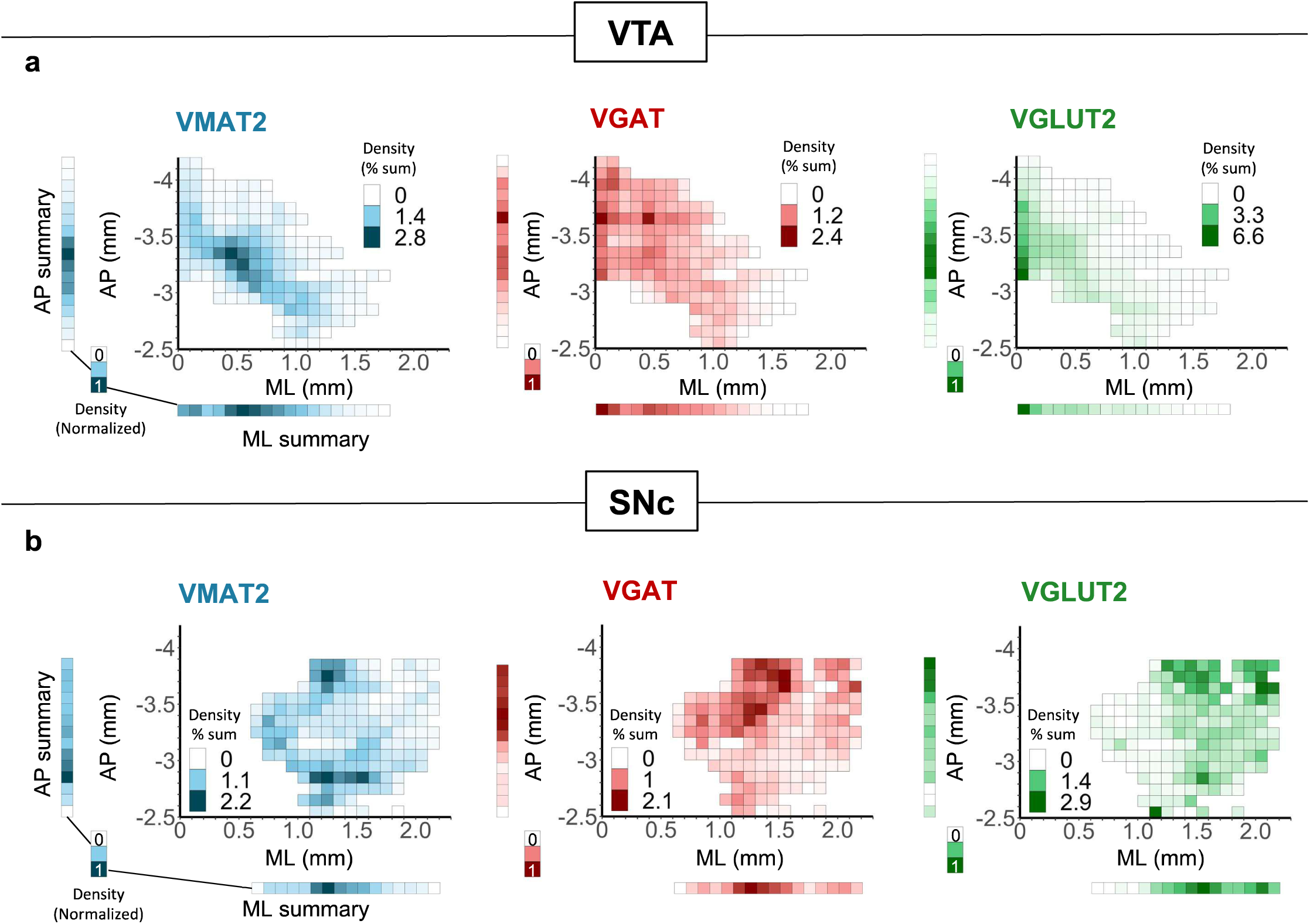
Spatial distribution of vesicular transporter-defined neurons along anterior- posterior (AP) and medial-lateral (ML) axes. (**A**) Density heatmaps of neurons expressing VMAT2, VGAT, or VGLUT2 across AP and ML axes. Summary bars display density data collapsed into either axis then normalized. (**B**) Same as (A) but for SNc.

For VTA (**Figure 7A**), VMAT2^+^ neuron density was centrally distributed with a hot spot around AP= -3.3, ML= 0.4. VGAT^+^ neurons are widely distributed but with a clear medial concentration and proportionally more representation in posterior areas compared to the other cell types. VGLUT2^+^ neurons were highly concentrated medially, and at any given ML coordinate can be seen shifted toward the anterior edge.

For SNc (**Figure 7B**), VMAT2^+^ neurons were densest along the ML extent at AP= -2.8 mm, with another hot spot at AP= -3.7, ML= 1.2. VGAT^+^ cells in SNc were most concentrated in medial-posterior SNc whereas VGLUT2^+^ cells were concentrated in lateral-posterior SNc.

Figure 7 plots all neurons that express a vesicular transporter and neurons may thus be represented in more than one group if they express more than one transporter. We made additional spatial heat maps for all 7 mutually exclusive cell types (**Supplemental** Figure 4). Consistent with prior reports, VGLUT2^+^ combinatorial cell types were medially concentrated in VTA but more laterally in SNc, consistent with the global pattern of VGLUT2 expression in these regions (Ma et al., 2023; Root et al., 2014; Steinkellner et al., 2018; Yamaguchi et al., 2013). In contrast, these maps reveal that while the global population of VGAT+ cells is most dense in posterior SNc and VTA, VGAT^+^ neurons that were also VMAT2^+^ were instead most prevalent in anterior SNc and VTA (**Supplemental** Figure 4B). Importantly, this finding is consistent with results from our reporter cross that also showed a pronounced reduction in the rate of TH colocalization with VGAT reporter in more posterior sections (**Supplemental** Figure 3B). Overall, these quantitative spatial analyses expand upon prior findings and highlight the distinct distribution pattern for each neurotransmitter-defined cell type within VTA and SNc.

## DISCUSSION

Prior works using IHC, ISH, scRNAseq or scPCR, and physiological assays have provided unequivocal evidence for the existence of DA, GABA, and glutamate-releasing neurons co-mingled in VTA and SNc (González-Hernández & Rodríguez, 2000; Kawano et al., 2006; Li et al., 2013; Mendez et al., 2008; Olson & Nestler, 2007; Phillips et al., 2022; Poulin et al., 2014; Tecuapetla et al., 2010; Tiklová et al., 2019; Tritsch et al., 2012; van Zessen et al., 2012; Yamaguchi et al., 2007, 2013, 2015). Previous works also established that these neuron types are not homogenously distributed within SNc and VTA, and demonstrated widespread neurotransmitter co-release from these overlapping neuronal populations (Reviewed in: Hnasko & Edwards, 2012; Morales & Margolis, 2017; Pupe & Wallén-Mackenzie, 2015; Trudeau et al., 2014). We undertook this study to more completely understand cellular diversity of these regions, as non-DA cell types also mediate functions relevant to addiction, Parkinson’s Disease, and related disorders. Indeed, DA modulation of dorsal striatal circuits are important for habitual and compulsive behaviors, (Reviewed in: Lerner, 2020; Luscher et al., 2020; O’Hare et al., 2018), while the role of SNc GABA- and glutamate-releasing neurons in this circuit have been little studied. Our results emphasize the prominence of non-DA cell types in VTA and SNc, which will be relevant for future work on diseases associated with mesolimbic and mesostriatal circuits.

Several studies have aimed to map the relative proportions of each of these transmitter-defined populations with variable levels of anatomical precision (Ma et al., 2023; Nair-Roberts et al., 2008; Simon et al., 2023). However, riboprobes used in some early ISH experiments likely had lower sensitivity, in part explaining why DA and other VTA neurons that express lower levels of VGLUT2 were undetected or underreported (Fremeau et al., 2001; Nair-Roberts et al., 2008; Yamaguchi et al., 2007). Based on these or other works, it is frequently asserted that only few glutamate neurons are present in SNc and VTA cell groups. This misconception is in stark contrast to many recent studies and the results reported here. Indeed, multiple studies highlight the functional relevance of VTA glutamate neurons in appetitive and aversive behaviors (Barbano et al., 2020; Qi et al., 2016; Warlow et al., 2023; Yoo et al., 2016; Zell et al., 2020), and endogenous VGLUT2 expression in SNc DA neurons confers resistance to neurotoxic insults (Buck et al., 2022; Dal Bo et al., 2008; Shen et al., 2018; Steinkellner et al., 2018, 2022).

IHC-based detection of glutamic acid decarboxylase (GAD1/2), VGAT, or VGLUT2 protein can be an unreliable approach to identify GABA or glutamate neurons in SNc and VTA. This is presumably due to a combination of factors, including comparatively lower levels of expression and the tendency for presynaptic proteins to concentrate at release sites rather than in neuronal soma (Fremeau et al., 2001; Hnasko & Edwards, 2012; Kaufman et al., 1991). For this reason, we relied on detection of their mRNA. Our approach using RNAscope to label mRNA encoding vesicular transporters offers several key advantages. The use of paired double “Z” probe hybridization results in low background, and when coupled with amplification offers high sensitivity (Wang et al., 2012), explaining why we detected more non-DA cells than reported in early ISH studies. Indeed, our findings are well aligned with recent studies that make use of genetic reporters or RNAscope. For example, recent findings have shown that VGLUT2 and VGAT drivers labeled similar numbers of cells in VTA (Root et al., 2020) and that neurons co-positive for TH and VGLUT2 represent a minority population (10-30%) of both the global DA and global glutamate neuron populations in VTA and SNc (Shen et al., 2018; Steinkellner et al., 2018, 2022; Warlow et al., 2023; Yan et al., 2018; Zell et al., 2020). Our findings also align well with scRNAseq studies that revealed an abundance of GABA and glutamate neurons, and identified subtypes of SNc and VTA neurons that go well beyond neurotransmitter markers (Azcorra et al., 2023; Phillips et al., 2022; Poulin et al., 2014; Tiklová et al., 2019; Viereckel et al., 2016). However, these studies cannot resolve subregional distribution and can be contaminated by dissections that include neighboring brain areas. Although patch-seq and laser capture microdissection approaches can circumvent this issue, they have been limited to small sample sizes and focused primarily on DA populations in VTA and SNc (Chung et al., 2005; Greene et al., 2005; Tapia et al., 2018; J. Wang et al., 2023). Our approach using RNAscope offers high signal-to-noise, high sampling rate, and high spatial resolution to extend prior findings.

Cellular neurotransmitter phenotypes are most typically determined by expression of genes required for neurotransmitter synthesis, recycling, and/or packaging into vesicles. Yet caution must be used when selecting genetic markers. For example, IHC against TH has been frequently used to reliably identify DA neurons, however not all cells expressing *Th* transcript have detectable TH protein or release detectable DA (Li et al., 2013; Stamatakis et al., 2013; Yamaguchi et al., 2015). Indeed, expression of the plasma membrane DA transporter (DAT, *Slc6a3*) can more reliably label neurons that express TH protein and is often used as a genetic driver because it is more selective for midbrain DA neurons than either TH or VMAT2, which are also expressed in other monoamine cell groups (Pupe & Wallén-Mackenzie, 2015). Further, there are multiple examples of a neuron releasing a neurotransmitter that it did not synthesize de novo, but instead recycled following its manufacture by another cell (Lebrand et al., 1996; Tritsch et al., 2014; Zhou et al., 2005). Because the synaptic release of dopamine, GABA, and glutamate ultimately relies on their packaging into vesicles, we chose to label the transcripts encoding the canonical vesicular transporters for the vesicular packaging and release of DA, GABA, and glutamate. However, even DA neurons that lack canonical markers for GABA synthesis and packaging were shown capable of releasing GABA in a process that depends on VMAT2 (Melani & Tritsch, 2022; Tritsch et al., 2012; Zych & Ford, 2022). Thus, describing VTA/SNc neurons as either a DA, GABA, or glutamate neuron based on expression of VMAT2, VGAT, VGLUT2, or any other marker is blurred by the capacity of subsets of these neurons to co-release multiple transmitters; a function enabled by expression of more than one of the canonical markers, and by non-canonical mechanisms for neurotransmitter synthesis, reuptake, and vesicular packaging.

We estimate that 20.4% of VTA and 10.4% of SNc neurons are positive for multiple vesicular transporters. Consistent with prior works, we observed a concentration of VMAT2^+^/VGLUT2^+^ cells in medial VTA and in lateral SNc (Hnasko et al., 2012; Kawano et al., 2006; Ma et al., 2023; Steinkellner et al., 2018; Yamaguchi et al., 2011). These findings are also consistent with observations that medial NAc is a hotspot for glutamate/dopamine co-transmission (Chuhma et al., 2014, 2023; Mingote et al., 2015), and that posterior striatum receives dense input from VGLUT2^+^ DA neurons (Azcorra et al., 2023; Poulin et al., 2018). Our results are also consistent with prior reports indicating a concentration of VGAT^+^/VGLUT2^+^ neurons in medial VTA (Root et al., 2018, 2020), and evidence for GABA/glutamate co-transmission in VTA projections to LHb and VP (Root et al., 2014; Yoo et al., 2016). In SNc, we also identified VGAT expression in 8.8% of VMAT2^+^ DA neurons. While at least some SNc DA neurons have been shown capable of releasing GABA from striatal terminals, this appears to depend on GABA packaging via VMAT2, as noted above. While it is unclear the extent to which VGAT might contribute to GABA release from a subset of DA neurons, our findings are consistent with recent scRNAseq and other data showing GABA markers are present in SNc DA neurons (Azcorra et al., 2023; Kim et al., 2015; Poulin et al., 2014; Saunders et al., 2018; Stamatakis et al., 2013; Tiklová et al., 2019). Finally, our identification of a small population of neurons that are positive for all three transporters is also consistent with other recent studies (Ma et al., 2023; Phillips et al., 2022), though whether such cells functionally release all three neurotransmitters remains an open question.

RNAscope-based ISH allows for exceptional signal to noise. However, because we report much higher levels of VGLUT2^+^ neurons in VTA/SNc than some early quantitative reports (Nair-Roberts et al., 2008; Yamaguchi et al., 2007), we ran additional experiments to validate our findings (see Figure 3). Based on these data it is unlikely that background signal is accounting for false positive detection. However there inevitably remains some uncertainty around the attribution of specific mRNA signals to specific cells, and this challenge is greater when cells are densely packed or when transcript levels within a cell are low. Because misattribution of signal could lead to false positives or false negatives when determining cellular rates of co-localization, we cross-validated our findings using a genetic reporter, used standardized attribution rules, and relied on two independent observers to minimize experimenter error (see methods). Importantly, our data are in broad agreement with other recent studies that have examined colocalization of one of these markers with another in SNc or VTA (Root et al., 2018, 2020; Shen et al., 2018; Steinkellner et al., 2018; Warlow et al., 2023; Yamaguchi et al., 2011; Yan et al., 2018; Zell et al., 2020).

While we argue that vesicular transporters are an equivalent or better marker of a neurotransmitter-defined neuron than alternatives, it is important to reiterate that a given vesicular transporter can package more than one transmitter. For example, VGAT can also package the inhibitory neurotransmitter glycine (Wojcik et al., 2006), though we are unaware of evidence that VGAT^+^ neurons in VTA/SNc release glycine. In addition to DA (and GABA), VMAT2 packages other monoamine neurotransmitters (Erickson et al., 1992; Peter et al., 1994). This includes serotonin-releasing neurons which are present in the most caudal areas of VTA as well as in the raphe nuclei caudal to VTA (Erickson et al., 1992; Weihe et al., 1994), somewhat compromising our ability to count DA neurons in the most caudal sections. Indeed, the frequency of *Tph2*^+^ neurons increased progressively in posterior sections through VTA and *Tph2* was found within VMAT2^+^ neurons in CLi, PBP, and IF. Based on analysis of 11 sections we estimate that 6.3% of VMAT2^+^ neurons in posterior region of VTA (Bregma -3.6 to -4.1) and 0.49% of total VMAT2^+^ VTA neurons that were included in our DA neuron counts, were instead *Tph*2^+^ serotonin neurons.

The anatomical borders between VTA, SNc, and neighboring regions are not always distinct (Swanson, 1982). Indeed, in caudal VTA we observed the intermixing of VTA DA and DR serotonin neurons (Figure 2). We also measured the highest number of VGLUT2^+^ neurons in anterior sections where subregions IF and RLi first appear (Figure 5A). Indeed, an abundance of VGLUT2^+^ neurons are present just anterior to these medial subregions, for example in supramammillary nucleus (Kesner et al., 2022). There is an analogous abundance of GABA neurons in structures posterior to VTA such as in RMTg (Jhou, 2005), or ventral to SNc in SNr. The extent to which such neighboring populations of glutamate and GABA neurons are genetically or functionally distinct from those that reside within the cytoarchitecturally defined boundaries of VTA and SNc remains an open and challenging question.

In sum, this report provides the most thorough anatomical description of neurotransmitter-defined cell types across VTA and SNc to date. Our observation that 78% of neurotransmitter-defined neurons in VTA express VGLUT2 and/or VGAT, as do 57% in SNc, emphasizes the enormous heterogeneity within these brain regions that are typically defined by their DA neurons. That significant proportions of each neurotransmitter-defined cell type express more than one vesicular transporter further highlights the diverse molecular landscape of these important brain areas. These data will thus serve as an important guide for future work unraveling the functions of these neurons.

## MATERIALS AND METHODS

### Animals

C57Bl/6J, *Slc17a6^tm2(cre)Lowl^* (VGLUT2-Cre), *Slc32a1^tm2(cre)Lowl^* (VGAT-Cre), and B6.Cg-Gt(ROSA)*^26Sortm6(CAG-ZsGreen1)Hze^*/J (ZsGreen) were obtained from The Jackson Laboratory (RRIDs: IMSR_JAX:000664, IMSR_JAX:016963, IMSR_JAX:016962, IMSR_JAX:007906) and bred in-house. Mice were bred at the University of California, San Diego (UCSD), group-housed (max 5 mice/cage), and maintained on a 12h light–dark cycle with food and water available *ad libitum*. All cages were equipped with bedding, cotton nesting material, and lofts. Mice were sacrificed at 8-10 weeks old for ISH experiment and 8-20 weeks old for combined IHC and ISH and VGAT reporter experiments. Male and female mice were included in all experiments, and all experiments were performed in accordance with protocols approved by the UCSD Institutional Animal Care and Use Committee.

### In situ hybridization (ISH)

Sample preparation and RNAscope labeling was followed as described (Conrad, 2024b). Briefly, experimenter’s gloves and tools were sprayed with RNAseZap (AM9780, ThermoFisher) prior to and between brain extractions. Mice were deeply anaesthetized with pentobarbital (200 mg/kg i.p.; Virbac) and decapitated. Brains were extracted, immediately snap frozen in isopentane chilled on dry ice, then stored at −80°C. Before sectioning, experimenter’s gloves and tools were again sprayed with RNAseZap. Brains were cut serially (20 μm) on a cryostat (CM3050S, Leica) as coronal sections through VTA and SNc. Sections were mounted directly onto SuperFrost Plus glass slides (Fisher), air-dried at room temperature (RT), then stored at - 20°C for 2 hours. Slides were stored long term at –80°C in an airtight container before starting the RNAscope assay (Advanced Cell Diagnostics, ACD fluorescent multiplex kit 320851). Briefly, slides were fixed in phosphate buffered saline (PBS) containing 4% paraformaldehyde at 4°C for 15 min then dehydrated in serial ethanol washes. Slides were incubated with protease IV at RT for 30 min. RNA hybridization was performed using antisense probes against *Slc17a6*/VGLUT2 (319171-C1), *Slc32a1*/VGAT (319191-C2), and *Slc18a2*/VMAT2 (425331-C3) for primary experiments. *Tph2* (ACD 318691), *Slc18a2*/VMAT2 (ACD 478351-C2), and *Th* (ACD 317621-C3) probes were used for Tph2 experiments. After the last wash step, slides were coverslipped using DAPI-containing Fluoromount-G (SouthernBiotech, 0100-01).

### Stereotaxic surgery

Surgery was conducted as described (Conrad & Oriol, 2024). Mice were anaesthetized with isoflurane, placed in a stereotaxic frame (Kopf), and injected bilaterally with AAV1-EF1a-FLEX- HTB (Salk Institute – GT3 core, 7x10^11 genomes/ml, RRID: Addgene_44187) into the VTA of homozygous VGLUT2-Cre mice (medial-lateral: ± 0.35, anterior-posterior: −3.4, dorsal-ventral: −4.4 mm relative to Bregma). 150 nl was injected per side at 10 nl/sec, 5 sec delay between 10 nl injection cycles (Nanoject III, Drummond Scientific) using pulled glass pipets (Drummond Scientific). For each infusion, the pipet was left in place for 10 min. The pipet was then raised 0.1 mm, left in place for 1 min, then slowly retracted. Animals were treated with the analgesic Carprofen (MWI, 5mg/kg s.c.) before and after surgery. Mice were monitored daily for the first five days and allowed to recover from surgery for 3 weeks, then sacrificed for the combined ISH and IHC assay.

### Combined in situ hybridization (ISH) and immunohistochemistry (IHC)

Dual RNAscope and IHC labeling was as described (Conrad, 2024c). In brief, these experiments used the antisense probe for Slc17a6/VGLUT2 (ACD 319171) and were conducted the same as ISH described above until the last wash step. After the last incubation step, slides were rinsed two times in PBS, blocked in PBS containing 4% normal donkey serum and 0.2% Triton X-100 for one hour at RT, then incubated with primary antibody (chicken Anti-GFP 1:10,000; Invitrogen, RRID: AB_2534023) overnight at 4°C in blocking buffer. The next day slides were washed for 5 min in PBS three times, then incubated with secondary antibody (anti-chicken 1:400; Alexa Fluor 488; Jackson ImmunoReseach, RRID: AB_2340376) for two hours at room temperature. Slides were washed for 15 min in PBS three times then briefly rinsed in wash buffer (ACD). Finally, slides were coverslipped using DAPI-containing Fluoromount-G.

### VGAT reporter IHC

Mice were deeply anaesthetized with pentobarbital (200 mg/kg i.p.; Virbac) and perfused using 4% PFA in PBS. The brains were stored in 4% PFA at 4 degrees overnight followed by 30% sucrose for 48 hours. The brains were then snap frozen in isopentane chilled on dry ice. Brains were cut serially (30 μm) on a cryostat (CM3050S, Leica) as coronal sections through VTA and SNc. Staining for TH was as described (Faget et al., 2024). Sections were rinsed two times in PBS, blocked in PBS containing 4% normal donkey serum and 0.2% Triton X-100 for one hour at RT, then incubated with primary antibody (Sheep Anti-TH 1:1,000; Pelfreez, RRID: AB_461070) overnight at 4°C in blocking buffer. The next day sections were washed for 5 min in PBS three times, then incubated with secondary antibody (anti-sheep 1:400; Alexa Fluor 647; Jackson ImmunoReseach, RRID: AB_2340751) for two hours at room temperature. Sections were washed for 15 min in PBS three times then received a 5 minute wash in DAPI (1:1,000; Sigma-Aldrich, D9542-10MG). Sectioned received a final PBS wash. Finally, slides were mounted and coverslipped using Fluoromount-G.

### Imaging

Image acquisition and processing for RNAscope experiments were followed as described (Conrad, 2024a). Tiled images were acquired using confocal microscopy (Zeiss LSM 880 Indimo, Zen Black 2, RRID:SCR_018163). All images were acquired using a 20X objective and focused with the Fluorescence Autofocus Strategy. Bit depth was set to 12, pixel averaging to 8, and pinhole AU to 1 on the blue channel. Each experiment type used different acquisition settings, but within an experiment type the acquisition settings were unchanged (**Supplemental Table 4**). Black value was adjusted in Zen for each image to minimize background signal, while white value was adjusted to maximize signal intensity near nuclei without saturation. For the VGAT reporter experiment, tiled images were acquired using a Zeiss Axio Observer.Z1. All images were acquired using a 20X objective and apotome. All images within this experiment were acquired with identical acquisition settings.

### Counting

For the triple transporter labeling experiment, sections covering the rostro-caudal extent of the VTA and SNc (-2.5 mm to -4.1 mm Bregma) at 100 µm intervals were included for cell quantification. A specific sample size was not calculated prior to experimentation, though we aimed to sample from each 100 um interval from at least one male and one female animal. We used 3 mice per sex and 3 to 10 sections per animal were used for analysis. The number of sections used for each Bregma point or subregion analysis is detailed in **Supplemental Table 3**. Scoring criteria and methods were as described (Conrad, 2024a). Briefly, isolated nuclei (circular DAPI signal with diameter ≥5 µm) were considered positive for RNA if at least 5 puncta were within ∼2 µm of nucleus edge and concentrically covering 50% or more of a nucleus’s circumference. In the case a nucleus touched neighboring nuclei, 30% of the RNA signal must unambiguously have been from the cell in question (i.e. signal cannot be readily ascribed to a neighboring cell). Neurons were marked positive using the event tool in Zen. Once neurons positive for individual mRNA were identified, we then looked for markers that colocalized to the same cell. Neurons were double checked to have both mRNA signals present, then were deemed co-positive. Each section was scored by two experimenters with concurrence of 93.1%, but neurons that were marked positive by only one observer were re-assessed before a final count was determined.

For the dual ISH and IHC experiment, any section with GFP expression in VTA (spanning -2.9 mm to -3.8 mm from bregma) was included and scored by a single experimenter. At least three sections per animal were used for analysis. A nucleus was considered GFP^+^ if GFP signal colocalized with DAPI signal. To assess false positive rate, we first drew boundaries enclosing GFP^+^ dense areas. Neurons were marked VGLUT2^+^ if they met the criteria described above, then assessed for GFP expression. To assess false negative rate, GFP^+^ nuclei were marked as having 0, 1, 2, 3, 4, or ≥5 puncta. The criteria of concentric puncta arrangement was excluded for this experiment since 1 punctum cannot be arranged in this manner.

For VGAT reporter experiment, counting was followed as described (Kollman & Hnasko, 2024). Nuclei were marked as ZsG positive if ZsG signal covered > 50% of the cell. If a cell was on the border of a region the cell was counted toward the region which contained at least 50% of the cell. Neurons were marked positive for ZsG using the event tool in Zen. Once neurons positive for ZsG were identified, we scored each cell for the presence or absence of TH colocalization to the same cell.

### Analysis

Cell counts were recorded in Excel workbooks (Microsoft, version 2403, RRID: SCR_016137) then visualized using RStudio (version 2022.07.2+576, RRID: SCR_000432) using base R (version 4.2.2, RRID: SCR_001905) with ggplot2 (3.4.0, RRID: SCR_014601), tidyverse (1.3.2, RRID: SCR_019186), and eulerr (6.1.1, RRID: SCR_022753) packages (Larsson, 2022; Wickham, 2016).

## AUTHOR CONTRIBUTIONS

William S. Conrad

Contribution: Conceptualization, Methodology, Validation, Formal Analysis, Investigation, Data Curation, Visualization, Writing – Original Draft Preparation, Reviewing, and Editing

Lucie Oriol

Contribution: Validation, Investigation, Writing – Review & Editing

Grace J. Kollman

Contribution: Validation, Investigation

Lauren Faget

Contribution: Validation, Writing – Review & Editing

Thomas S. Hnasko

Contribution: Conceptualization, Methodology, Validation, Resources, Supervision, Project Administration, Funding Acquisition, Writing – Original Draft Preparation, Reviewing and Editing

## ACKNOWLEDGEMENTS

For the purpose of open access, the author has applied a CC-BY 4.0 public copyright license to the Author Accepted Manuscript (AAM) version arising from this submission.

## DATA AND MATERIAL AVAILABILITY

For the triple labeling experiment, the related images, cell count data sheets, and code for visualization are available at Zenodo repository (10.5281/zenodo.10607238). Tabular data for all figures is available at the following repository (10.5281/zenodo.11189934).

