## Supplemental Figures and Tables for "Proportion and distribution of neurotransmitter-defined cell types in the ventral tegmental area and substantia nigra pars compacta"

| Bregma (mm) | TPH2+ neuron counts | VMAT2+ neuron counts | Percentage of VMAT2+ neurons positive for TPH2 | Mean percent for bregma point |
|---|---|---|---|---|
| **-3.6** | 2 | 135 | 1.5 | 3.1 |
| | 2 | 127 | 1.6 | |
| | 7 | 94 | 7.4 | |
| **-3.7** | 13 | 109 | 11.9 | 9.0 |
| | 4 | 80 | 5 | |
| **-3.9** | 4 | 73 | 8.2 | 8.2 |
| **-4.0** | 2 | 24 | 8.3 | 7.0 |
| | 2 | 33 | 6.1 | |
| **-4.1** | 4 | 16 | 25 | 29.6 |
| | 4 | 11 | 36.4 | |
| **-4.2** | 13 | 27 | 48.1 | 48.1 |

**Supplemental Table 1 (related to Figure 2). Fraction of TPH2/VMAT2 copositive neurons in caudal VTA (unilateral).**

| Bregma (mm) | Region | ZsG+ neuron counts | ZsG+ neurons with TH | Percent of co-positive neurons |
|---|---|---|---|---|
| -3.0 | VTA | 146 | 46 | 31.5 |
| -3.1 | VTA | 182 | 31 | 17 |
| -3.2 | VTA | 221 | 45 | 20.4 |
| -3.4 | VTA | 187 | 31 | 16.6 |
| -3.4 | VTA | 111 | 15 | 13.5 |
| -3.5 | VTA | 131 | 17 | 13 |
| -3.7 | VTA | 154 | 4 | 2.6 |
| -3.7 | VTA | 153 | 0 | 0 |
| -3.8 | VTA | 146 | 2 | 1.4 |
| -3.9 | VTA | 122 | 4 | 3.3 |
| -3.9 | VTA | 110 | 0 | 0 |
| -4.1 | VTA | 17 | 1 | 5.9 |
| -3.0 | SNc | 76 | 30 | 39.5 |
| -3.1 | SNc | 79 | 18 | 22.8 |
| -3.2 | SNc | 101 | 20 | 19.8 |
| -3.4 | SNc | 82 | 17 | 20.7 |
| -3.4 | SNc | 138 | 21 | 15.2 |
| -3.5 | SNc | 146 | 18 | 12.3 |
| -3.7 | SNc | 123 | 18 | 14.6 |
| -3.7 | SNc | 113 | 6 | 5.3 |
| -3.8 | SNc | 92 | 7 | 7.6 |
| -3.9 | SNc | 110 | 5 | 4.5 |
| -3.9 | SNc | 34 | 1 | 2.9 |

**Supplemental Table 2 (related to Figure 4, Supplemental Figure 3). Fraction of ZsG/TH copositive neurons in VTA and SNc of VGAT-Cre reporter mice.**

| | | n (mice) | n (sections) | Sex | VMAT2 only | VGAT only | VGLUT2 only | VMAT2$^+$/ VGAT$^+$ | VMAT2$^+$/ VGLUT2$^+$ | VGAT$^+$/ VGLUT2$^+$ | Triple | Total |
|---|---|---|---|---|---|---|---|---|---|---|---|---|
| VTA global | Cumulative cell count | 3 | 25 | M | 2597 | 2417 | 1906 | 166 | 871 | 771 | 139 | 8867 |
| | | 3 | 23 | F | 2716 | 1871 | 1865 | 129 | 585 | 654 | 102 | 7922 |
| | | 6 | 48 | B | 5313 | 4288 | 3771 | 295 | 1456 | 1425 | 241 | 16789 |
| | Fraction (%) | 3 | 25 | M | 29.3 | 27.3 | 21.5 | 1.9 | 9.8 | 8.7 | 1.6 | - |
| | | 3 | 23 | F | 34.3 | 23.6 | 23.5 | 1.6 | 7.4 | 8.3 | 1.3 | - |
| | | 6 | 48 | B | 31.6 | 25.5 | 22.5 | 1.8 | 8.7 | 8.5 | 1.4 | - |
| PIF | Cumulative cell count | 2 | 7 | M | 221 | 63 | 60 | 2 | 41 | 24 | 5 | 416 |
| | | 2 | 7 | F | 199 | 46 | 121 | 4 | 24 | 20 | 2 | 416 |
| | | 4 | 14 | B | 420 | 109 | 181 | 6 | 65 | 44 | 7 | 832 |
| | Fraction (%) | 2 | 7 | M | 53.1 | 15.1 | 14.4 | 0.5 | 9.9 | 5.8 | 1.2 | - |
| | | 2 | 7 | F | 47.8 | 11.1 | 29.1 | 1.0 | 5.8 | 4.8 | 0.5 | - |
| | | 4 | 14 | B | 50.5 | 13.1 | 21.8 | 0.7 | 7.8 | 5.3 | 0.8 | - |
| PN | Cumulative cell count | 2 | 9 | M | 254 | 211 | 176 | 10 | 173 | 26 | 19 | 869 |
| | | 2 | 9 | F | 324 | 150 | 171 | 8 | 124 | 36 | 5 | 818 |
| | | 4 | 18 | B | 578 | 361 | 347 | 18 | 297 | 62 | 24 | 1687 |
| | Fraction (%) | 2 | 9 | M | 29.2 | 24.3 | 20.3 | 1.2 | 19.9 | 3.0 | 2.2 | - |
| | | 2 | 9 | F | 39.6 | 18.3 | 20.9 | 1.0 | 15.2 | 4.4 | 0.6 | - |
| | | 4 | 18 | B | 34.3 | 21.4 | 20.6 | 1.1 | 17.6 | 3.7 | 1.4 | - |
| PBP | Cumulative cell count | 3 | 22 | M | 1824 | 1552 | 678 | 135 | 387 | 321 | 63 | 4960 |
| | | 3 | 21 | F | 1965 | 1332 | 679 | 102 | 267 | 282 | 54 | 4681 |
| | | 6 | 43 | B | 3789 | 2884 | 1357 | 237 | 654 | 603 | 117 | 9641 |
| | Fraction (%) | 3 | 22 | M | 36.8 | 31.3 | 13.7 | 2.7 | 7.8 | 6.5 | 1.3 | - |
| | | 3 | 21 | F | 42.0 | 28.5 | 14.5 | 2.2 | 5.7 | 6.0 | 1.2 | - |
| | | 6 | 43 | B | 39.3 | 29.9 | 14.1 | 2.5 | 6.8 | 6.3 | 1.2 | - |
| RLi | Cumulative cell count | 3 | 13 | M | 49 | 47 | 548 | 4 | 55 | 220 | 17 | 940 |
| | | 2 | 11 | F | 8 | 36 | 473 | 2 | 26 | 138 | 10 | 693 |
| | | 5 | 24 | B | 57 | 83 | 1021 | 6 | 81 | 358 | 27 | 1633 |
| | Fraction (%) | 3 | 13 | M | 5.2 | 5.0 | 58.3 | 0.4 | 5.9 | 23.4 | 1.8 | - |
| | | 2 | 11 | F | 1.2 | 5.2 | 68.3 | 0.3 | 3.8 | 19.9 | 1.4 | - |
| | | 5 | 24 | B | 3.5 | 5.1 | 62.5 | 0.4 | 5.0 | 21.9 | 1.7 | - |
| IF | Cumulative cell count | 2 | 10 | M | 58 | 39 | 197 | 5 | 81 | 98 | 22 | 500 |
| | | 2 | 10 | F | 88 | 70 | 199 | 8 | 66 | 127 | 23 | 581 |
| | | 4 | 20 | B | 146 | 109 | 396 | 13 | 147 | 225 | 45 | 1081 |
| | Fraction (%) | 2 | 10 | M | 11.6 | 7.8 | 39.4 | 1.0 | 16.2 | 19.6 | 4.4 | - |
| | | 2 | 10 | F | 15.1 | 12.0 | 34.3 | 1.4 | 11.4 | 21.9 | 4.0 | - |
| | | 4 | 20 | B | 13.5 | 10.1 | 36.6 | 1.2 | 13.6 | 20.8 | 4.2 | - |
| VTAR | Cumulative cell count | 3 | 6 | M | 95 | 77 | 104 | 4 | 12 | 45 | 0 | 337 |
| | | 2 | 7 | F | 86 | 45 | 146 | 1 | 14 | 36 | 1 | 329 |
| | | 5 | 13 | B | 181 | 122 | 250 | 5 | 26 | 81 | 1 | 666 |
| | Fraction (%) | 3 | 6 | M | 28.2 | 22.8 | 30.9 | 1.2 | 3.6 | 13.4 | 0 | - |
| | | 2 | 7 | F | 26.1 | 13.7 | 44.4 | 0.3 | 4.3 | 10.9 | 0.3 | - |
| | | 5 | 13 | B | 27.2 | 18.3 | 37.5 | 0.8 | 3.9 | 12.2 | 0.2 | - |
| CLi | Cumulative cell count | 3 | 8 | M | 96 | 428 | 143 | 6 | 122 | 37 | 13 | 845 |
| | | 2 | 5 | F | 46 | 192 | 76 | 4 | 64 | 15 | 7 | 404 |
| | | 5 | 13 | B | 142 | 620 | 219 | 10 | 186 | 52 | 20 | 1249 |
| | Fraction (%) | 3 | 8 | M | 11.4 | 50.7 | 16.9 | 0.7 | 14.4 | 4.4 | 1.5 | - |
| | | 2 | 5 | F | 11.4 | 47.5 | 18.8 | 1.0 | 15.8 | 3.7 | 1.7 | - |
| | | 5 | 13 | B | 11.4 | 49.6 | 17.5 | 0.8 | 14.9 | 4.2 | 1.6 | - |
| SNc global | Cumulative cell count | 3 | 22 | M | 1845 | 1681 | 412 | 179 | 219 | 22 | 25 | 4383 |
| | | 3 | 21 | F | 1749 | 1319 | 345 | 158 | 212 | 16 | 28 | 3827 |
| | | 6 | 43 | B | 3594 | 3000 | 757 | 337 | 431 | 38 | 53 | 8210 |
| | Fraction (%) | 3 | 22 | M | 42.1 | 38.4 | 9.4 | 4.1 | 5.0 | 0.5 | 0.6 | - |
| | | 3 | 21 | F | 45.7 | 34.5 | 9.0 | 4.1 | 5.5 | 0.4 | 0.7 | - |
| | | 6 | 43 | B | 43.8 | 36.5 | 9.2 | 4.1 | 5.2 | 0.5 | 0.6 | - |
| SNcD | Cumulative cell count | 3 | 21 | M | 1133 | 938 | 221 | 150 | 130 | 11 | 4 | 2587 |
| | | 2 | 20 | F | 1126 | 705 | 230 | 128 | 156 | 10 | 11 | 2366 |
| | | 5 | 41 | B | 2259 | 1643 | 451 | 278 | 286 | 21 | 15 | 4953 |
| | Fraction (%) | 3 | 21 | M | 43.8 | 36.3 | 8.5 | 5.8 | 5.0 | 0.4 | 0.2 | - |
| | | 2 | 20 | F | 47.6 | 29.8 | 9.7 | 5.4 | 6.6 | 0.4 | 0.5 | - |
| | | 5 | 41 | B | 45.6 | 33.2 | 9.1 | 5.6 | 5.8 | 0.4 | 0.3 | - |
| SNcL | Cumulative cell count | 3 | 18 | M | 82 | 330 | 171 | 9 | 64 | 11 | 21 | 688 |
| | | 3 | 16 | F | 59 | 246 | 89 | 8 | 47 | 6 | 17 | 472 |
| | | 6 | 34 | B | 141 | 576 | 260 | 17 | 111 | 17 | 38 | 1160 |
| | Fraction (%) | 3 | 18 | M | 11.9 | 48.0 | 24.9 | 1.3 | 9.3 | 1.6 | 3.1 | - |
| | | 3 | 16 | F | 12.5 | 52.1 | 18.9 | 1.7 | 10.0 | 1.3 | 3.6 | - |
| | | 6 | 34 | B | 12.2 | 49.7 | 22.4 | 1.5 | 9.6 | 1.5 | 3.3 | - |
| SNcM | Cumulative cell count | 3 | 15 | M | 571 | 317 | 18 | 19 | 12 | 0 | 0 | 937 |
| | | 2 | 13 | F | 510 | 245 | 26 | 19 | 6 | 0 | 0 | 806 |
| | | 5 | 28 | B | 1081 | 562 | 44 | 38 | 18 | 0 | 0 | 1743 |
| | Fraction (%) | 3 | 15 | M | 60.9 | 33.8 | 1.9 | 2.0 | 1.3 | 0 | 0 | - |
| | | 2 | 13 | F | 63.3 | 30.4 | 3.2 | 2.4 | 0.7 | 0 | 0 | - |
| | | 5 | 28 | B | 62.0 | 32.2 | 2.5 | 2.2 | 1.0 | 0 | 0 | - |
| SNcV | Cumulative cell count | 3 | 9 | M | 59 | 96 | 2 | 1 | 13 | 0 | 0 | 171 |
| | | 2 | 9 | F | 54 | 123 | 0 | 3 | 3 | 0 | 0 | 183 |
| | | 5 | 18 | B | 113 | 219 | 2 | 4 | 16 | 0 | 0 | 354 |
| | Fraction (%) | 3 | 9 | M | 34.5 | 56.1 | 1.2 | 0.6 | 7.6 | 0 | 0 | - |
| | | 2 | 9 | F | 29.5 | 67.2 | 0.0 | 1.6 | 1.6 | 0 | 0 | - |
| | | 5 | 18 | B | 31.9 | 61.9 | 0.6 | 1.1 | 4.5 | 0 | 0 | - |

**Supplemental Table 3 (related to Figures 4 & 5). Cell-type counts and fractions for subregions disaggregated by sex.**

| Experiment | Label | Laser λ (nm) | Laser source | Laser Power | Gain (ms) |
|---|---|---|---|---|---|
| ***Slc17a6/Slc32a1/Slc18a2*** | DAPI | 405 | Diode | 0.50% | 611 |
| | *Slc17a6* | 488 | Argon | 0.50% | 558 |
| | *Slc32a1* | 561 | Diode-pumped solid-state | 0.50% | 750 |
| | *Slc18a2* | 633 | Helium-Neon | 0.50% | 700 |
| **Dual ISH and IHC** | DAPI | 405 | Diode | 0.50% | 611 |
| | GFP | 488 | Argon | 0.15% | 558 |
| | *Slc17a6* | 561 | Diode-pumped solid-state | 0.50% | 750 |
| **TPH2** | DAPI | 405 | Diode | 0.50% | 611 |
| | *Slc18a2* | 488 | Argon | 0.40% | 558 |
| | *Tph2* | 561 | Diode-pumped solid-state | 0.18% | 750 |
| | *Th* | 633 | Helium-Neon | 0.50% | 700 |

**Supplemental Table 4 (related to methods). Image acquisition settings.**

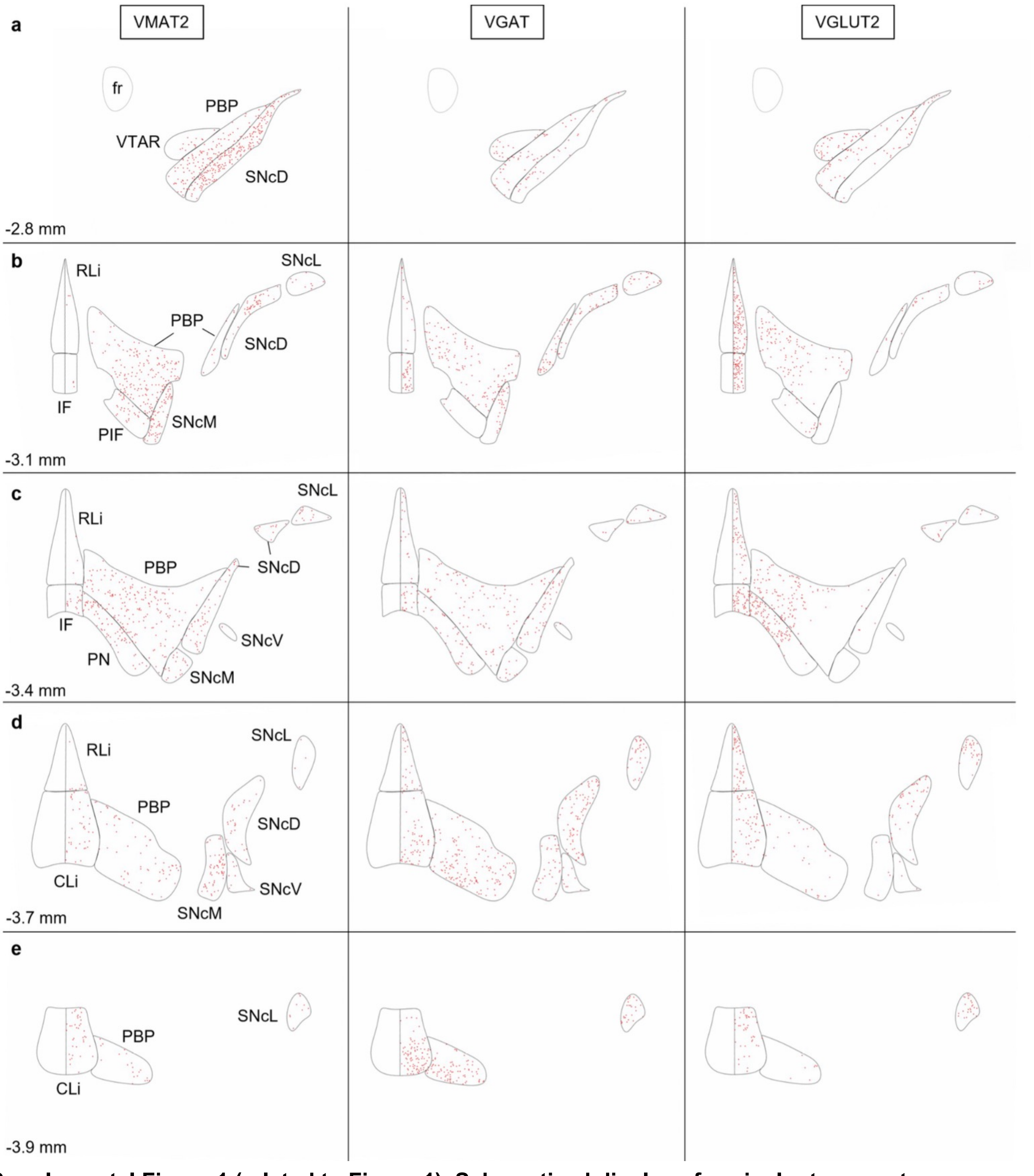


**Supplemental Figure 1 (related to Figure 1). Schematized display of vesicular transporter-expressing neurons.** (**A-E**) Each plot is from the corresponding image in **Figure 1** and each red dot represents the location of a counted VMAT2$^{+}$, VGAT$^{+}$, or VGLUT2$^{+}$ neuron. CLi (caudal linear nucleus), fr (fasciculus retroflexus), IF (interfascicular nucleus), PBP (parabrachial pigmented nucleus), PIF (parainterfascicular nucleus), PN (paranigral nucleus), RLi (rostral nucleus), SNcD (SNc dorsal), SNcL (SNc lateral), SNcM (SNc medial), SNcV (SNc ventral), VTAR (VTA, rostral).

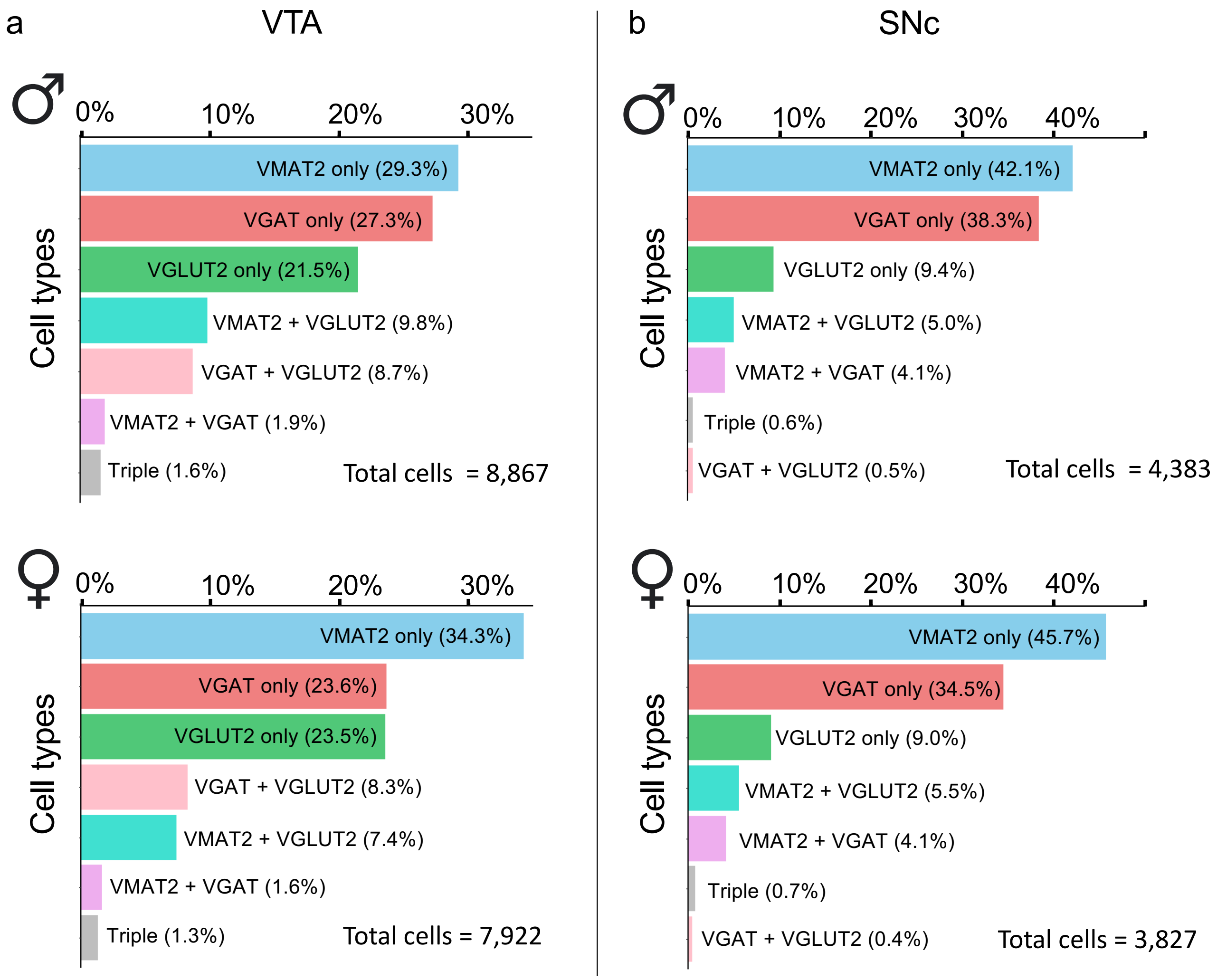


**Supplemental Figure 2 (related to Figure 4). Global proportions of vesicular transporter-defined neurons in VTA and SNc displayed by sex. (A)** Fraction of labeled VTA neurons that expressed one or more vesicular transporter in VTA of male (top panel) and female (bottom panel) mice. (**B**) Same as (A) but for SNc.

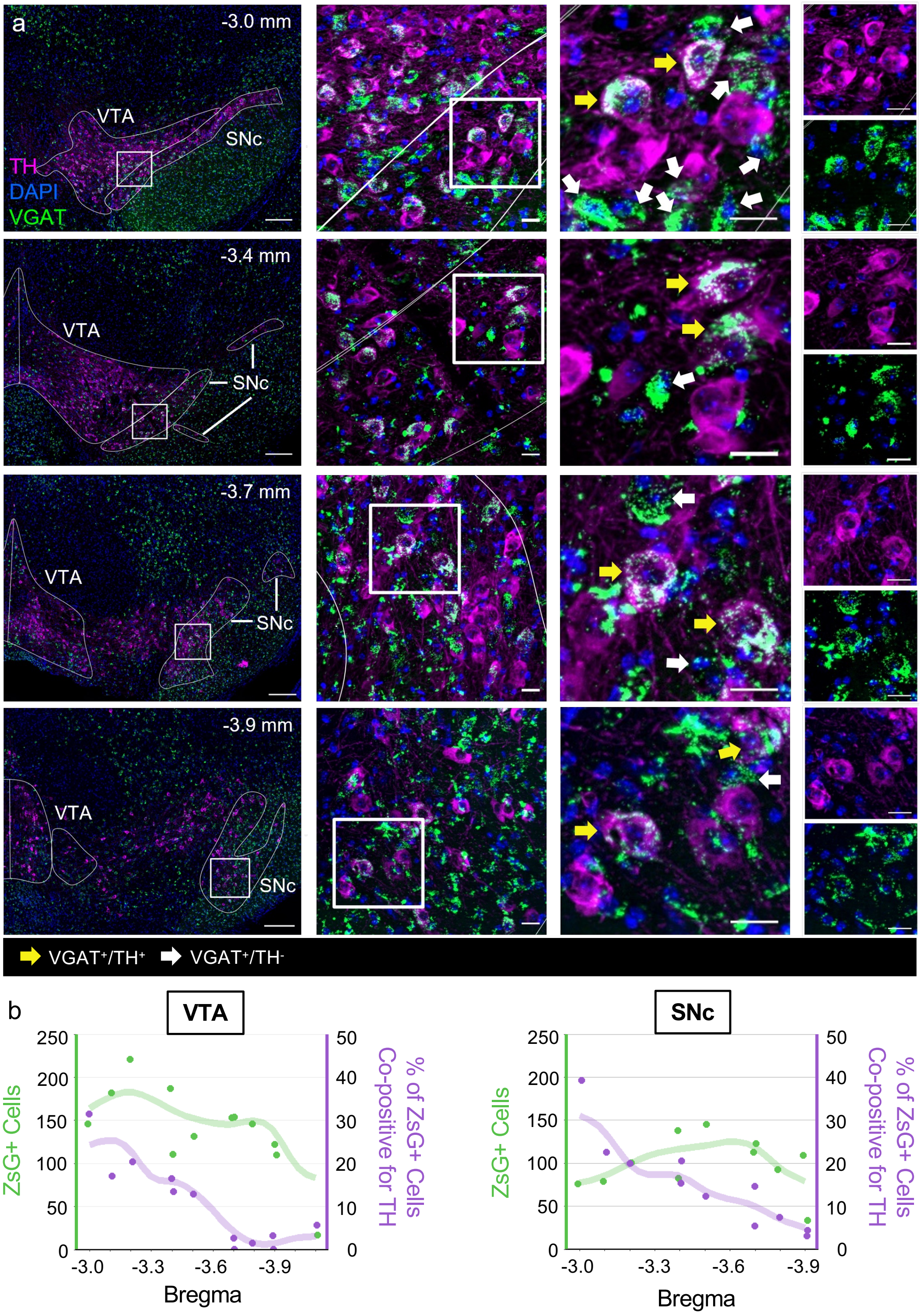


**Supplemental Figure 3 (related to Figure 4). ZsG/TH co-positive neurons in VTA and SNc of VGAT-Cre reporter mice. (A)** Example images of coronal sections from VGAT-Cre x ZsGreen reporter (green) mice immunostained for TH (magenta), with DAPI (indigo). Left panels show wide-field view with VTA and SNc demarcated, scale 200 μM. Middle left panels (scale, 20 μM) represent white box from left panel; middle right panels (scale, 20 μM) represent white box from middle left panel. Yellow arrows point to neurons co-positive for ZsG and TH, white arrows point to ZsG+ neurons negative for TH. Right panels separately show TH (top) and ZsG (bottom) signal from middle right panels. **(B)** Cell counts for ZsG+ cells and percentage of ZsG+ cells co-positive for TH by Bregma. Trend lines are moving averages smoothed with spline regression.

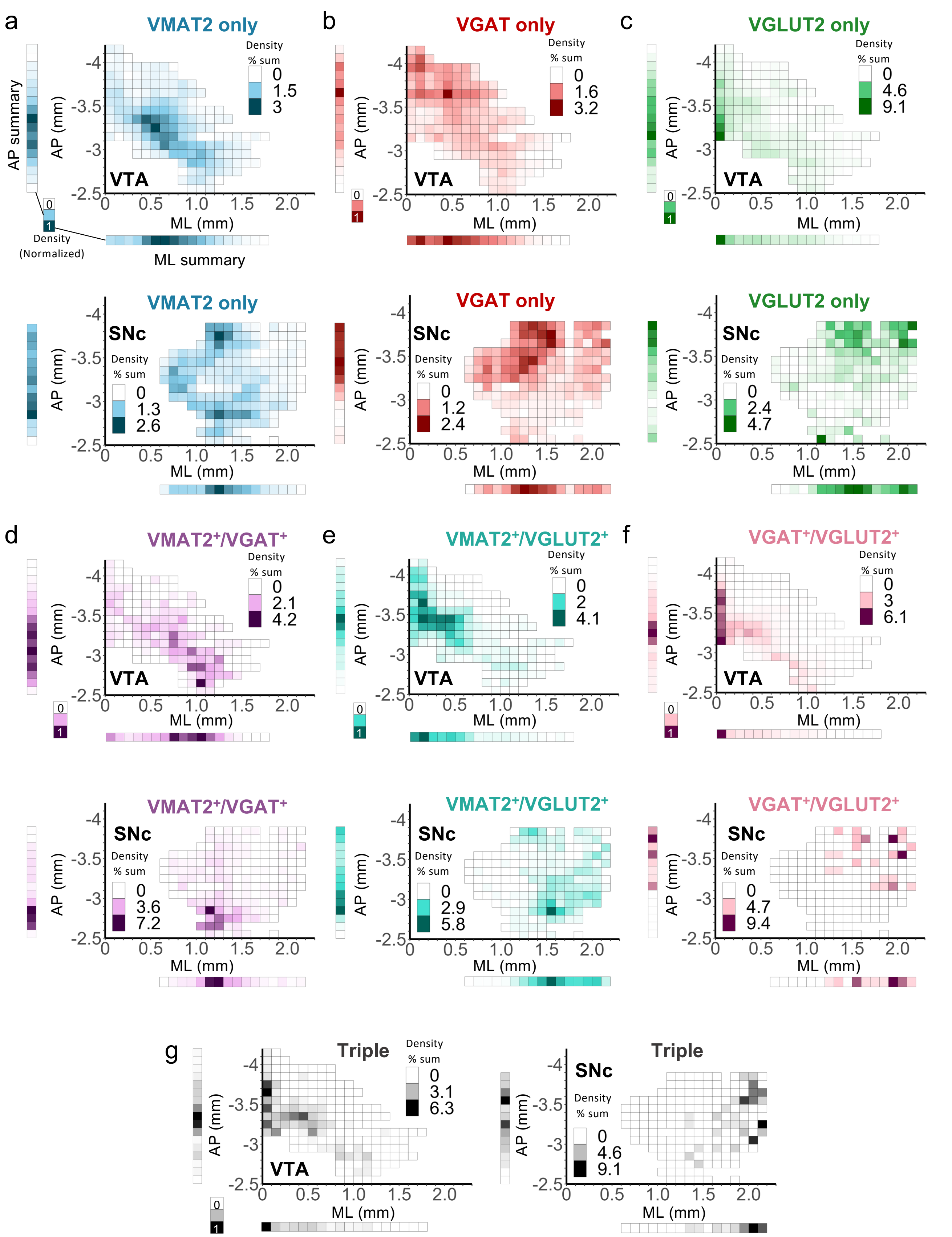


**Supplemental Figure 4 (related to Figure 6). Spatial distribution of vesicular transporter-defined neurons along anterior-posterior (AP) and medial-lateral (ML) axes.** Density heatmaps of neurons expressing (**A**) VMAT2-only, (**B**) VGAT-only, (**C**) VGLUT2-only, (**D**) VMAT2$^{+}$/VGAT$^{+}$, (**E**) VMAT2$^{+}$/VGLUT2$^{+}$, (**F**) VGAT$^{+}$/VGLUT2$^{+}$ or (**G**) all three vesicular transporters (Triple) across AP and ML axes. Summary bars display density data collapsed into either axis then normalized.
